# Hierarchical Generalized Linear Mixed Model for Genome-wide Association Analysis

**DOI:** 10.1101/2021.03.10.434742

**Authors:** Hengyu Zhang, Li’ang Yang, Yanan Xu, Xiaojing Zhou, Yuxin Song, Shuling Li, Runqing Yang

## Abstract

In genome-wide association analysis (GWAS) for binary traits, we stratified the genomic generalized linear mixed model (GLMM) into two hierarchies—the GLMM regarding genomic breeding values (GBVs) and a generalized linear regression of the normally distributed GBVs to the tested marker effects. In the first hierarchy, the GBVs were predicted by solving for the genomic best linear unbiased prediction for GLMM with the estimated variance components or genomic heritability in advance, and in the second hierarchy, association tests were performed using the generalized least square (GLS) method for the GBVs. Like the Hi-LMM for regular quantitative traits, the so-called Hi-GLMM method exhibited higher statistical power to detect quantitative trait nucleotides (QTNs) with better genomic control for complex population structure than existing methods, especially when the GBVs were estimated precisely and using joint association analysis for QTN candidates obtained from a test at once. Application of the Hi-GLMM to re-analyze maize kernel colors and six human diseases illustrated its advantage over existing GLMM-based association methods in terms of computing efficiency and statistical power.

## Introduction

Using random polygenic effects excluding the tested marker to correct confounders such as population stratification and cryptic relatedness, the linear mixed model (LMM) can efficiently control the false positive rate and improve the power to detect quantitative trait nucleotides (QTNs). However, because of the requirement of the high computing intensity for LMM, researchers were motivated to develop simplified algorithms (Aulchenko, et al., 2007; Kang, et al., 2010; Kang, et al., 2008; Lippert, et al., 2011; Loh, et al., 2015; Svishcheva, et al., 2012; Zhang, et al., 2010; Zhou and Stephens, 2012). In general, LMM, which assumes the phenotype is distributed normally, is appropriate for continuous quantitative traits (Henderson, 1984). For complex disease traits expressed as a binary phenotype, however, such a genome-wide mixed model association does not provide interpretable and predictable mapping results (Yang, et al., 2014).

As a type of quantitative traits, complex diseases are thought to be controlled by a number of loci each having a small effect on the phenotype (Bulmer, 1971; Falconer, 1981). Instead of the linear regression model, logistic regression in the generalized linear model (GLM) (McCullagh and Nelder, 1989; Wedderburn, 1974) has also been applied to analyze the association between markers of interest in binary disease phenotypes. Despite the correction for fixed-effect covariates (Joel and Witte, 2012; Zaitlen, et al., 2012; Zaitlen, et al., 2012), logistic regression still produces inflation of association test statistics. With consideration of random polygenic effects, the generalized linear mixed model (GLMM) (Breslow and Clayton, 1993) was consequently proposed to decrease the false positive rate in mapping QTNs for disease traits. However, genome-wide GLMM-based association requires much more computation time than the mixed model association with either restricted maximum likelihood estimation (REML) (Schall, 1991) or Markov chain Monte Carlo (MCMC) iteration (Sorenrsen and Gianola, 2002). In addition, when the maximum likelihood for estimation and approximations are used to avoid numerical integration, GLMM leads to serious bias induced by the approximations (Gilmour, et al., 1985), and solutions tend to the positive/negative infinity.

In ascertained case-control studies when the proportions of cases and controls are not a random sample from the population, GLMM gives biased estimates of genomic hereditability for disease traits (Lee, et al., 2011) and therefore suffers from a loss in power to detect QTNs. Based on the calibrated genomic hereditability for case-control ascertainment, a Chi-squared score statistic for GWAS of disease traits was computed from posterior mean liabilities (PMLs) under the liability-threshold model (Hayeck, et al., 2015). To simplify the GLMM-based association analysis, the GMMAT (Chen, et al., 2016) and SAIGE (Zhou, et al., 2018) separately extend the EMMAX (Kang, et al., 2010) and BOLT-LMM (Loh, et al., 2015) from normally distributed traits to binary diseases. Using null model to association tests, GMMAT can detect QTNs for binary traits in high computing efficiency, but accurately estimate QTN effects. Moreover, SAIGE may produce false positive error caused by complex relationship structure in animal and plant breeding population, as the BOLT-LMM proposed for relatively independent human population. By means of sparse matrix (GRM)-based algorithms (Jiang, et al., 2021), SAIGE was applied into fast dissect binary diseases in the large-scale UK Biobank data, without well controlling false positive errors.

In this study, we partitioned the genomic GLMM into two hierarchies by defining genomic breeding values (GBVs) subjected to normal distribution for liability. The GBVs could be predicted by solving for the genomic best linear unbiased prediction (GBLUP) for GLMM in the first hierarchy, and then in the second hierarchy, association tests were performed by generalized linear regression of the estimated GBVs to the SNP effect tested. As an extension of Hi-LMM (Hao, et al., 2021) to binary traits, the hierarchical GLMM association analysis is called Hi-GLMM in short. The utility of Hi-GLMM is illustrated by extensive computer simulations and real data analyses.

## Methods

### Hi-GLMM

For binary disease traits, a logit regression model based on binomial distribution defines the linear relationship between phenotypes of the traits and the genetic effect of the SNPs tested. To govern the false positive rate for mapping QTNs, random polygenic effects as the confounding variables are considered as additional predictors. Thus, the genomic logit mixed model (Breslow and Clayton, 1993) was constructed as follows:0

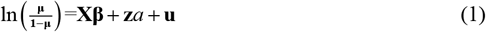

where **μ** is population mean of binary disease traits, **β** is a vector of fixed non-genetic effects; a is additive genetic effect of the tested SNP; **X** is the incidence matrix for **β**; **z** is a vector of indicator variables of the SNP genotypes, and **u** is a vector of random polygenic effects excluding the SNP tested, which is assumed to have normal distribution 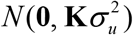 with an unknown polygenic variance 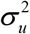 and a genomic relationship matrix (GRM) **K** (VanRaden, 2008) generally calculated from the entire genomic markers.

In linear predictors of GLMM (1), we define GBVs as

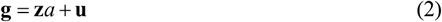

Then, GLMM (1) can be divided into two hierarchies:

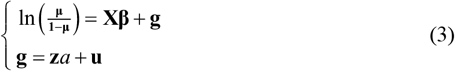

At this moment, **u** is regarded as the residuals in the model at the second hierarchy.

The model at the first hierarchy is a GLMM regarding fixed non-genetic effects and random GBVs. In the GLMM, both genomic heritability and continuous GBVs can be estimated with penalized quasi-likelihood for the GLMM (Breslow and Clayton, 1993; Chen, et al., 2016; Zhou, et al., 2018). Whereas, the model at the second hierarchy is a linear regression model for the GBVs normally distributed. Due to the heterogeneous polygenic covariance matrices, the genetic effect of the tested SNPs should be statistically inferred by the generalized least square (GLS) (Kariya and Kurata, 2004).

Such a hierarchical solution for genome-wide GLMM associations is summarized as follows:

1) Estimate GBVs with GBLUP for the Logit mixed model. Based on the GLMM at the first hierarchy, we construct the following GBLUP

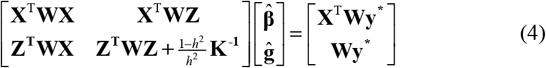

with **W** = **μ**(**1** − **μ**) and 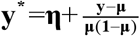 Where, **y** is a binary phenotype, 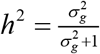 is the genomic heritability with a GBV variance of 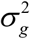 and residual variance of 1 which is assumed in GLMM.
2) Estimate genetic effect of the tested SNP with GLS By using diagonal eigenvalue matrix **D** and eigenvector matrix **U** from the eigen-decomposed **K** = **UDU**^T^, we spectrally transformed **ĝ**, **z** and **u** to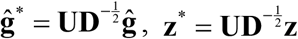 and 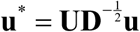. The model at the second hierarchy becomes 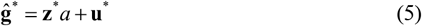 According to 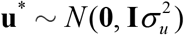, we estimate genetic effect

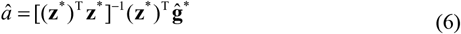

and its variance of genetic effect

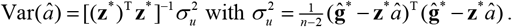
3) Statistically infer QTN by the statistic: 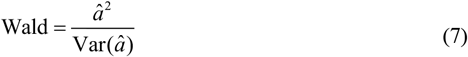 Which subject to Chi-squared distribution with 1 degree of freedom.
4) Joint association analysis

After one test at a time, some SNPs are chosen as QTN candidates at the significance level lower than the stringent Bonferroni corrected criterion (Hochberg and Tamhane, 1987). It should be noted that number of QTN candidates are generally limited to less than the population size in solving multiple linear regressions. With consideration of possible linkage disequilibrium among QTN candidates, we jointly analyzed multiple QTN candidates to improve the statistical power to detect QTNs in the following linear regression model:

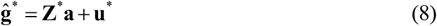

Where, **Z**\***a** are the regression terms of QTN candidates. Given the significance level of 0.01, the genetic effects are selected stepwise with backward regression approach, and at Bonferroni corrected criterion, the corresponding QTNs can also be identified according to the corrected statistic (7).

Based on the Hi-LMM, GBVs for liability are estimated with GBLUP for null GLMM, and a user-friendly Hi-GLMM software with the options of a test at once and joint association analysis was developed, which is freely available at https://github.com/RunKingProgram/Hi-GLMM.

### Simulated phenotypes

Two genomic datasets in maize (Romay, et al., 2013) and human (Wellcome Trust Case Control, 2007) were used to simulate the adaptability of our proposed Hi-GLMM to population structure. The maize population had more complex structure than the human population. We extracted 300,000 SNPs for both 2640 maize and 5004 human samples through higher quality control. In whole simulations, phenotypic incidences were all controlled at 0.5. QTNs were distributed randomly over the entire SNPs, and the additive effects were sampled from a gamma distribution with shape = 1.66 and scale = 0.4. Given the genomic heritability of liability, phenotypes of the control (0) and case (1) can be generated from the genomic logit model (1).

In addition to the population structure, the number of QTNs, genome heritability, and sampling number of SNPs were considered as experimental factors in the simulations. Under almost perfect genomic control, the ROC profiles were plotted by statistical powers to detect QTNs relative to a given series of Type I errors. Statistical powers are defined as the percentage of identified QTNs that have the maximum test statistic among their 20 closest neighbors over the total number of simulated QTNs. Simulations were repeated 50 times and the simulated positions and effects of QTNs varied in each simulation. The average results were recorded.

### Real phenotypes

The two datasets were used to illustrate the performance of Hi-GLMM: (1) total 2,631 inbred lines have been genotyped at 681,258 SNPs in maize, and of which 1595 recorded for kernel colors (KC) (Romay, et al., 2013). (2) in the Wellcome trust case-control consortium (WTCCC) study 1 (Wellcome Trust Case Control, 2007), there were the 3,004 shared controls and 11,985 cases from six common diseases, such as bipolar disorder (BD), coronary artery disease (CAD), rheumatoid arthritis (RA), hypertension (HT), type I diabetes (T1D) and type II diabetes (T2D). A total of 490,032 SNPs were genotyped in the human population.

## Results

### Statistical properties of the Hi-GLMM

For the two genomic datasets, we simulated 40, 200, and 1000 QTNs at the low (0.2), moderate (0.5), and high (0.8) genomic heritability, respectively, to control the phenotypes. The Hi-GLMM, a test at once was comparable with the four competing methods: GRAMMAR for binary traits (Song, et al., 2022), GMMAT, LTMLM, and SAIGE. The Q-Q and ROC profiles are displayed in Figure 1 and Figure 2 for the simulated phenotypes controlled by 200 QTNs, respectively, and are provided in Figure 1S and Figure 2S for all the simulated phenotypes. We also recorded genomic control values for each method in Table 1S. Under ideal genomic control very close to 1, Hi-GLMM behaves stably and exhibits a slightly higher statistical power than the GMMAT which approximates the exact GLMM, irrespective of how much QTNs control the quantitative traits and how complex the population structures are. GRAMMAR produces the highest false negative errors among Hi-GLMM and the four competing methods, and the population structure is more complex, its false negative rate is larger. For the simulated phenotypes in maize, LTMLM was found to be superior to the other methods in terms of statistical power, but it produced strong false positive errors. Similarly, SAIGE also exhibited higher statistical power with false positive errors for those controlled by 1,000 QTNs at the genomic heritability of 0.2. For the human population, there was no distinct difference in statistical properties between Hi-GLMM and the four competing methods. GRAMMAR yielded some false negative errors.

**Table 1.**
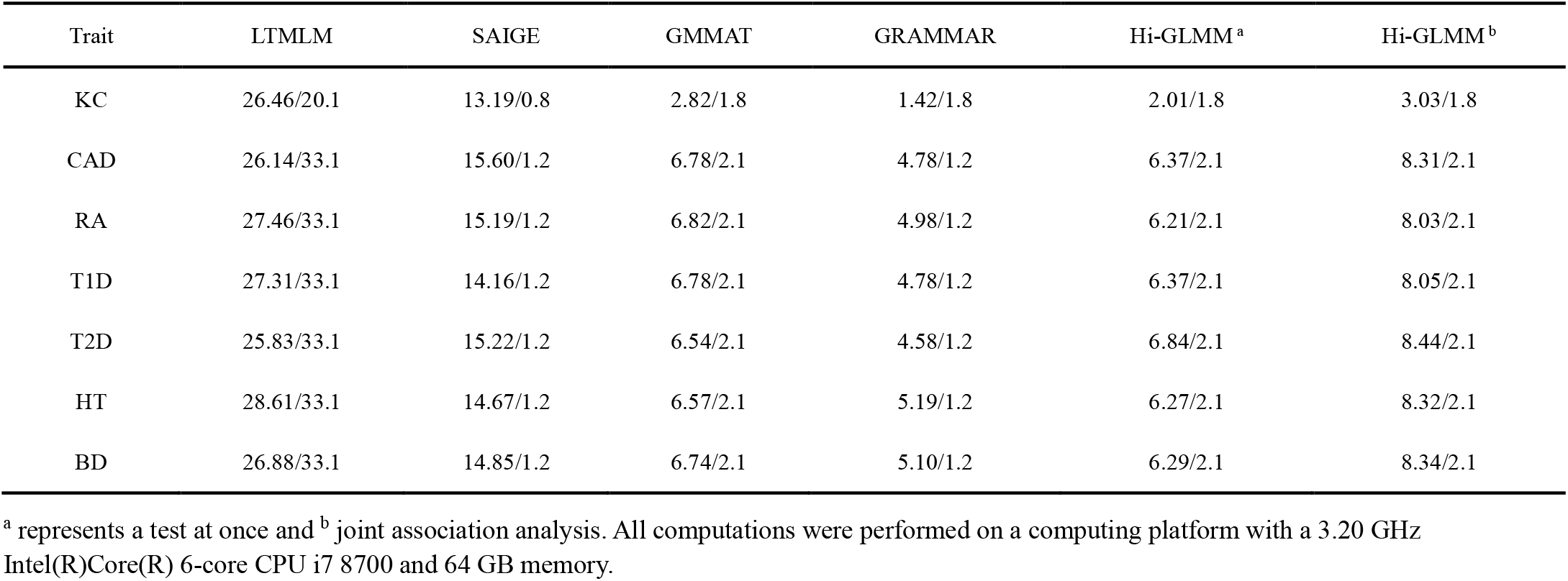
Computing time/memory footprints (Min/Gb) of the four competing methods for the six diseases.

**Figure 1:**
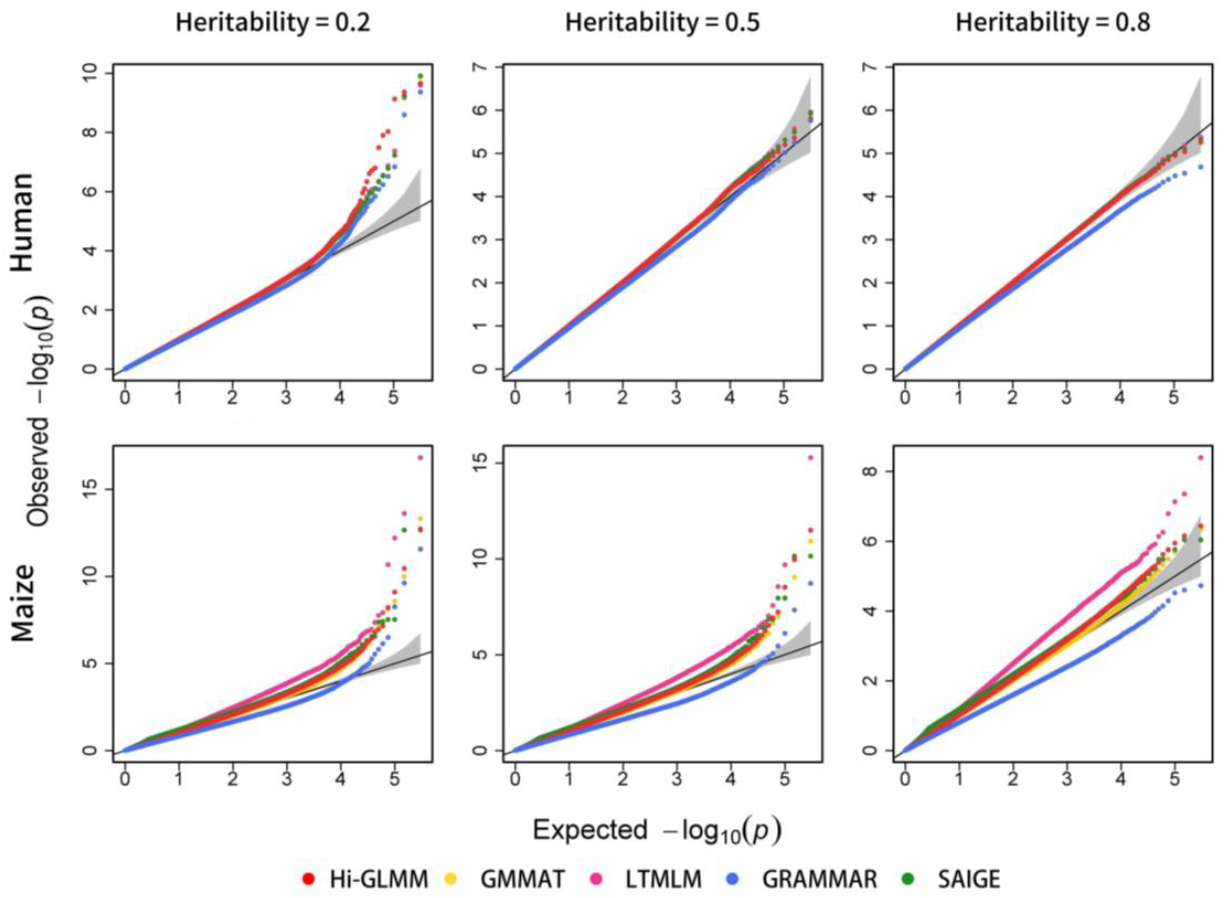
Comparison of Hi-GLMM with the four competing methods in the Q-Q profiles. The simulated phenotypes are controlled by 200 QTNs with the low, moderate and high heritabilities in maize and human populations. The Q-Q profiles for all simulated phenotypes are reported in Supplementary Figure 1S.

**Figure 2:**
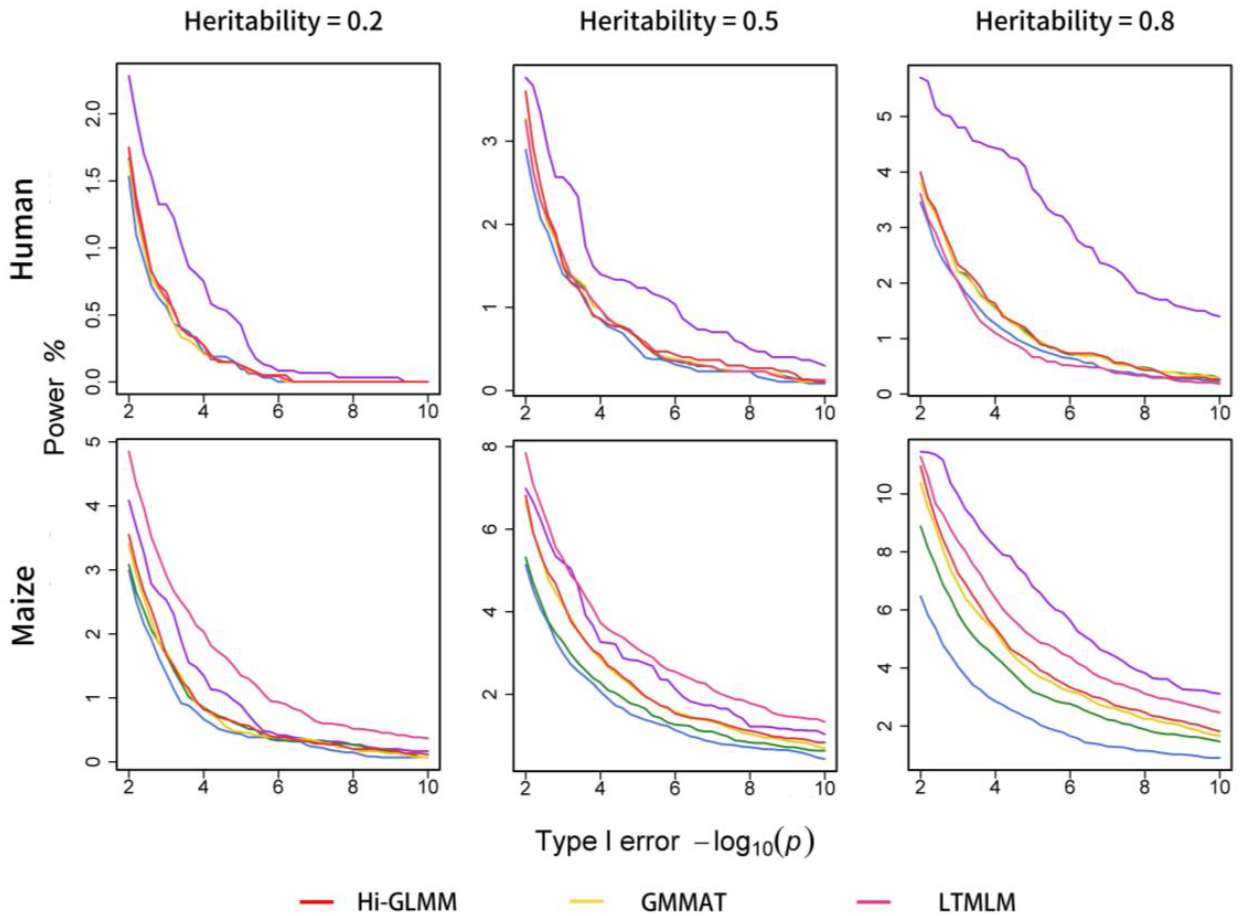
Comparison of Hi-GLMM with the four competing methods in the ROC profiles. The ROC profiles are plotted using the statistical powers to detect QTNs relative to the given series of Type I errors. Here, the simulated phenotypes are controlled by 200 QTNs with the low, moderate and high heritabilities in maize and human populations. The ROC profiles for all simulated phenotypes are reported in Supplementary Figure 2S.

Furthermore, we jointly analyzed multiple QTN candidates chosen from the Hi-GLMM, a test at once at the significance level of 0.05. For convenience of comparison, the statistical powers obtained with joint association analyses were depicted along with those using a test at once. Through backward multiple regression analysis, joint association analyses exhibited significantly the improved statistical power in almost the same genomic control as a test at once. Even though producing the highest false positive rates, LTMLM with a test at once was inferior to Hi-GLMM with joint analysis in the terms of statistical power.

### Sensitivity to estimate genomic heritability or GBVs

When genomic heritability or GBVs were precisely estimated, the Hi-LMM achieved much higher statistical power to detect QTNs, than EMMAX and BOLT-LMM. To test whether this finding fits Hi-GLMM, GMMAT, and SAIGE, which are extensions of Hi-LMM, EMMAX, and BOLT-LMM, respectively, for binary traits, we analyzed the simulated phenotypes controlled by 200 QTNs at the different heritabilities by taking the simulated genomic heritability or GBVs as the estimated polygenic effects. As shown in Figure 3 and Figure 3S, for the Hi-GLMM, one test at a time exhibited higher statistical power with the more ideal genomic controls than joint association analysis, if genomic heritability or breeding values were full estimated accurately. In contrast, GMMAT showed a somewhat reduced both statistical power and genomic control. Especially, SAIGE did not find any QTNs that were removed from the residual phenotypes. In addition, we adopted a Lasso technique implemented in R/glmnet (Friedman, et al., 2010) to precisely estimate the GBVs and found that Hi-GLMM produced higher statistical power than GMMAT using the improved the GBVs. This suggested that Hi-GLMM was expected to boost statistical power by more accurately estimating genomic heritability or GBVs.

**Figure 3:**
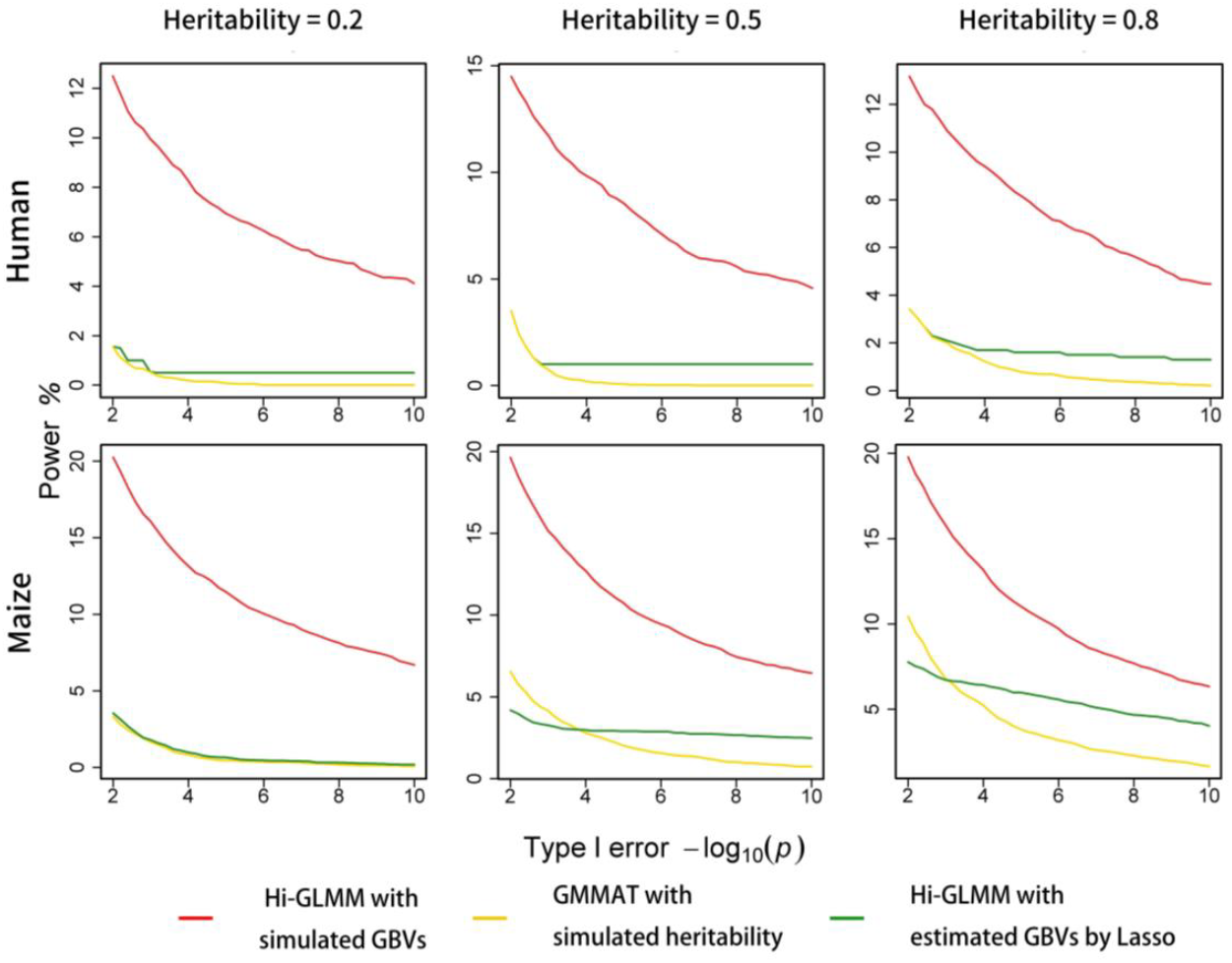
Sensitivity of statistical powers to estimate heritabilities or GBVs for Hi-GLMM. Statistical powers are dynamically evaluated with the ROC profiles. Both GRAMMAR and SAIGE do not detect any QTN with the simulated GBVs. The simulated phenotypes are controlled by 200 QTNs with the low, moderate and high heritabilities in maize and human populations.

### Calculation of GRMs with sampling markers

Using sampling markers to estimate the GRMs, we improved computational efficiency of the Hi-LMM. To test whether this finding fits Hi-GLMM, we randomly took 10 K, 20 K, 30 K, 40 K, 50K and 60K SNPs from the entire genomic markers and analyzed the phenotypes controlled by different numbers of QTNs at the heritability of 0.5 with all the methods, except LTMLM in which SNPs can be not sampled to estimate heritability. Figure 4 shows changes in the genomic controls with sampling levels of SNPs. Hi-GLMM gradually controlled positive false errors and improved statistical power to detect QTNs (Figure 4S and Figure 5S) with the sampling markers increased, similar to the results of GMMAT. In addition, Hi-GLMM needed more sampling markers than GMMAT to yield the ideal genomic control and statistical power using all genomic markers, but GMMAT often made genomic control unstable at larger sampling markers. In contrast, SAIGE produced serious false negative errors in the complex maize population irrespective of how many SNPs were used.

**Figure 4:**
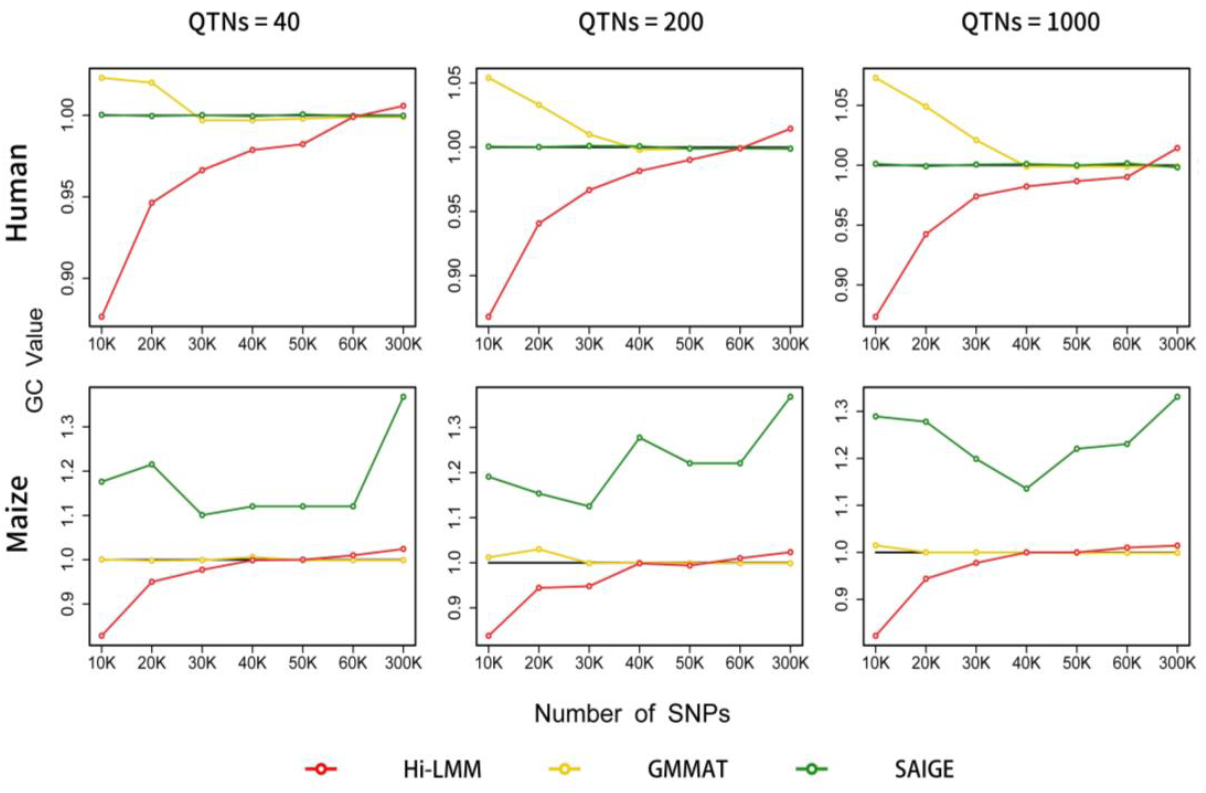
Changes in genomic controls with the number of sampling SNPs for Hi-GLMM and the five competing methods. Genomic control is calculated by averaging genome-wide test statistics. The simulated phenotypes are controlled by 40, 200 and 1000 QTNs with the moderate heritability in maize and human populations.

### Real data analysis

For the two genomic datasets, a standard quality control (QC) was conducted to exclude SNPs with MAFs < 0.01 and HWE > 0.05, and remove individuals with missing rates > 0.01. After the QC process, about 550,000 and 410,000 SNPs in maize and human were separately remained for the generalized mixed model association analyses. There were the incidences of 19.70% for kernel colors, while 39.82%-40.00% for six diseases.

In maize, genomic heritability for kernel colors were estimated to be 0.87 with penalized quasi-likelihood method. we analyzed the binary traits with the four competing methods and depicted the Q-Q and Manhattan profiles in Figure 7S. GMMAT and SAIGE detected almost the same 7 QTNs for kernel colors on Chromosomes 1, 2, 6, 7 and 9. In comparison, GRAMMAR searched the least 3 QTNs in higher false negative error and LTLMM did the most those with the highest false positive error. With one test at once, the Hi-GLMM found only 3 QTNs on Chromosomes 1, 3 and 6, but total 8 QTNs were detected on Chromosomes 1, 2, 3, 6 and 9 via joint association analysis. Different from the four competing methods, the Hi-GLMM did not detect any significant SNPs on Chromosome 7.

For the six diseases, we estimate genomic heritability to be 0.856, 0.976, 0.777, 0.931, 0.819 and 0. 872, respectively, for BD, CAD, HT, RA, T1D and T2D with penalized quasi-likelihood method. The Q-Q and Manhattan profiles for the six common diseases are shown in Figure 6S which indicate results obtained by Hi-GLMM and the four competing methods used in simulations. We concluded that (1) under perfect genomic control, Hi-GLMM found the QTNs for each disease on each chromosome, and the numbers of the detected QTNs were not less than those obtained by all the competing methods; and (2) in Hi-GLMM, joint association analyses detected more QTNs than one test at once. In particular, GRAMMAR detected least QTNs with the lowest genomic control among all the methods. GMMAT found more SNPs whose −log(p) exceeded the Bonferroni corrected thresholds for CAD, T1D, T2D, and HT with the highest false positive errors. Additionally, LTMLM abnormally estimated genomic heritabilities with scale transformation (Lee, et al., 2011), making genomic control for CAD, BD, T2D and HT unstable.

We conducted all data analyses on a computing platform with a 3.20 GHz Intel(R)Core(R) 6-core CPU i7 8700 and 64 GB memory. For real phenotypes, the running time and memory footprints were recorded from input of genotypes and phenotypes to the output of resulting QTNs for all methods. Table 1 showed that Hi-GLMM took less computing time than LTLMM and SAIGE. GMMAT consumed almost the same computing time and memory footprint as Hi-GLMM, but could not accurately estimate genetic effects. GRAMMAR performed higher computing efficiency with lower memory footprints than Hi-GLMM, but produced high false negative errors in detecting QTNs.

## Discussion

The GLMM formulates the relationship between binary response variables and linear predictors for liability through link function (McCullagh and Nelder, 1989; Wedderburn, 1974). As random linear predictors, the GBVs are estimated as a normal variable by the GBLUP equations for GLMM (Breslow and Clayton, 1993). In association tests in the second hierarchy, this allows Hi-GLMM to statistically infer the QTNs for binary traits with the GLS for the normally distributed GBVs successfully obtained with the GLMM in the first hierarchy. For the same independent variables, linear models are superior to generalized linear models on computing efficiency. While genomic heritability and GBVs can be well estimated, the Hi-GLMM will achieve genome-wide association tests faster, as compared to the GMMAT and SAIGE based on generalized linear model (Chen, et al., 2016; Zhou, et al., 2018).

In the LTMLM, genome-wide GLMM-based association analysis was also stratified into two hierarchies: first estimating PMLs by Bayesian sampling under the liability-threshold model (Hayeck, et al., 2015), and then, statistically inferring the QTNs with the LMM for the PMLs. However, both Bayesian sampling and LMM association analysis for PMLs showed that the LTMLM produced a high computing cost and serious false positive errors under complex structured populations, as illustrated in simulations. Like the GMMAT and SAIGE, the Hi-GLMM had to estimate the genomic heritability and GBVs using genomic markers. In contrast, the famers over-estimated polygenic heritabilities and effects by genomic heritability and GBVs, potentially increasing false negative rates, while the later unbiasedly and best estimated polygenic effects with generalized linear regression at the second hierarchy, ensuring high statistical power to detect QTNs in good genomic control.

Genome-wide GLMM association analysis is being devoted to improve statistical power and handle a large-scale population. Within the framework of Hi-GLMM, precise estimation for GBVs will contribute to attain high statistical power to detect QTNs, but this depends on the development of genomic selection for binary traits (Gianola, 2013). Furthermore, joint association analysis for multiple QTN candidates obtained with a test at once significantly improved the statistical power because possible linkage disequilibrium among candidate markers is considered in stepwise regression.

To speed up the Hi-GLMM, the GRM can be re-constructed with markers selected randomly across the whole genome (Jennifer, et al., 2012). Theoretically, the GRM may be better estimated with less and relatively independent markers separately drawn from all linkage groups than those randomly drawn from the whole genome, but construction of linkage groups may additionally increase the computational cost. In addition, transformation of human GRM to be the sparse (Jiang, et al., 2021) can also greatly simplify the Hi-GLMM to analyze a large-scale population, as long as genomic heritability or GBVs are well estimated.

## Competing interests

The authors declared that they have no conflicts of interest to this work.

## Author contribution statement

RQY proposed Hi-GLMM method. HYZ, LAY and YXS wrote the source code and data analysis, YNX, XJZ and SLL collected real datasets and participated in interpreting results. RQY, HYZ and SLL wrote this manuscript. All authors read and approved the final manuscript.

## Data availability

The datasets can be downloaded freely from https://www.panzea.org/%21#genotypes/cctl for phenotypic and genomic data in maize, and http://www.wtccc.org.uk/ with authorization for human genomic data.

## Fundings

Heilongjiang Bayi Agricultural University Support Program for San Heng San Zong (ZRCPY202024) and the National Natural Science Foundations of China (32072726).

## Notes

### Competing Interest Statement

The authors have declared no competing interest.

### Summary of Updates

Author affiliations updated; Methods have been theoretically corrected; A mew dataset has analyzed and added in sections of real phenotypes and Results; Supplemental files updated.

