## Supplementary material for "Hierarchical Generalized Linear Mixed Model for Genome-wide Association Analysis": Figure1S-7S and Table 1S

**Figure 1S:** Comparison in the Q-Q profiles between Hi-GLMM and the four competing methods. The simulated phenotypes are controlled by 40, 200, 1,000 QTNs with the low, moderate and high heritabilities in 1) human and 2) maize.

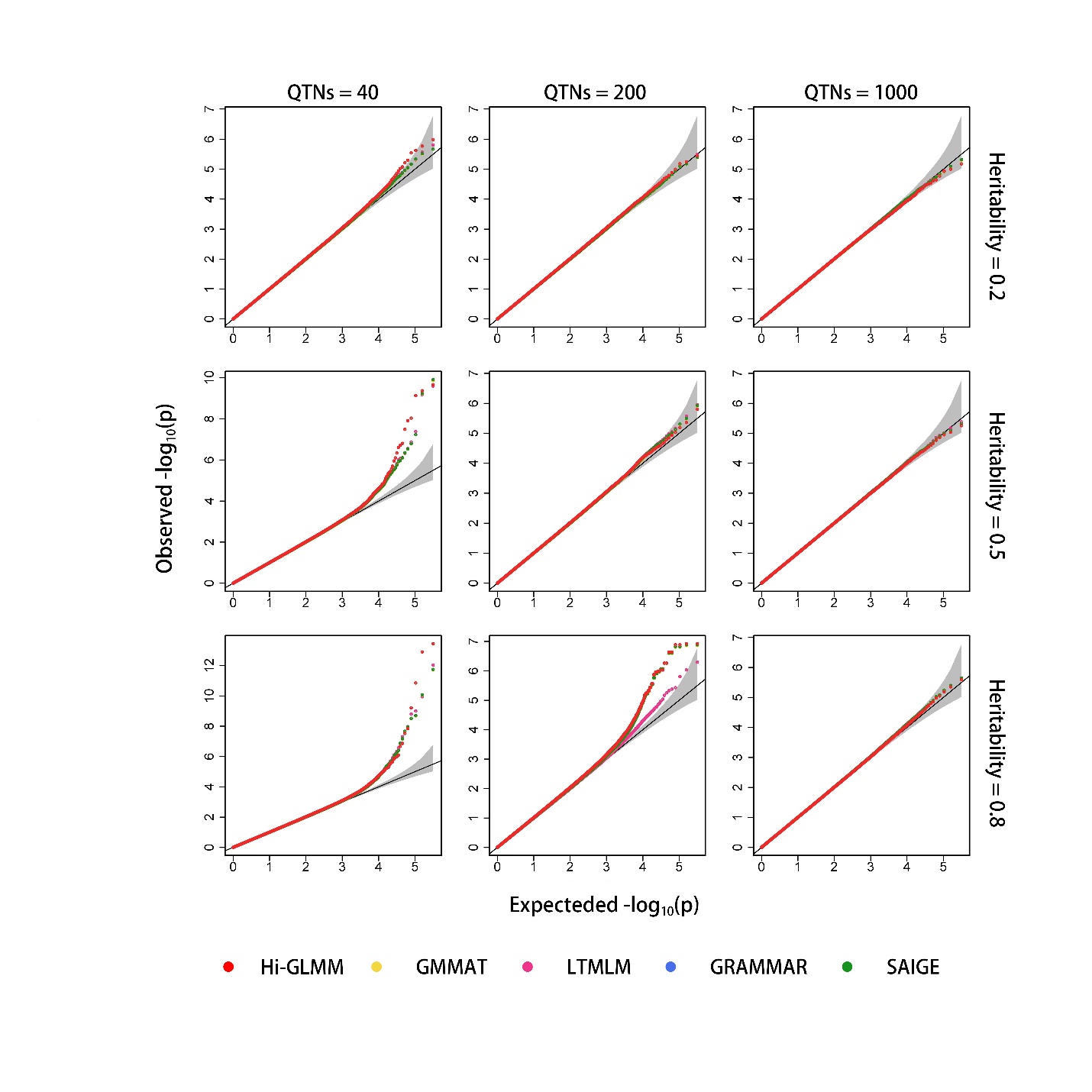

(1)

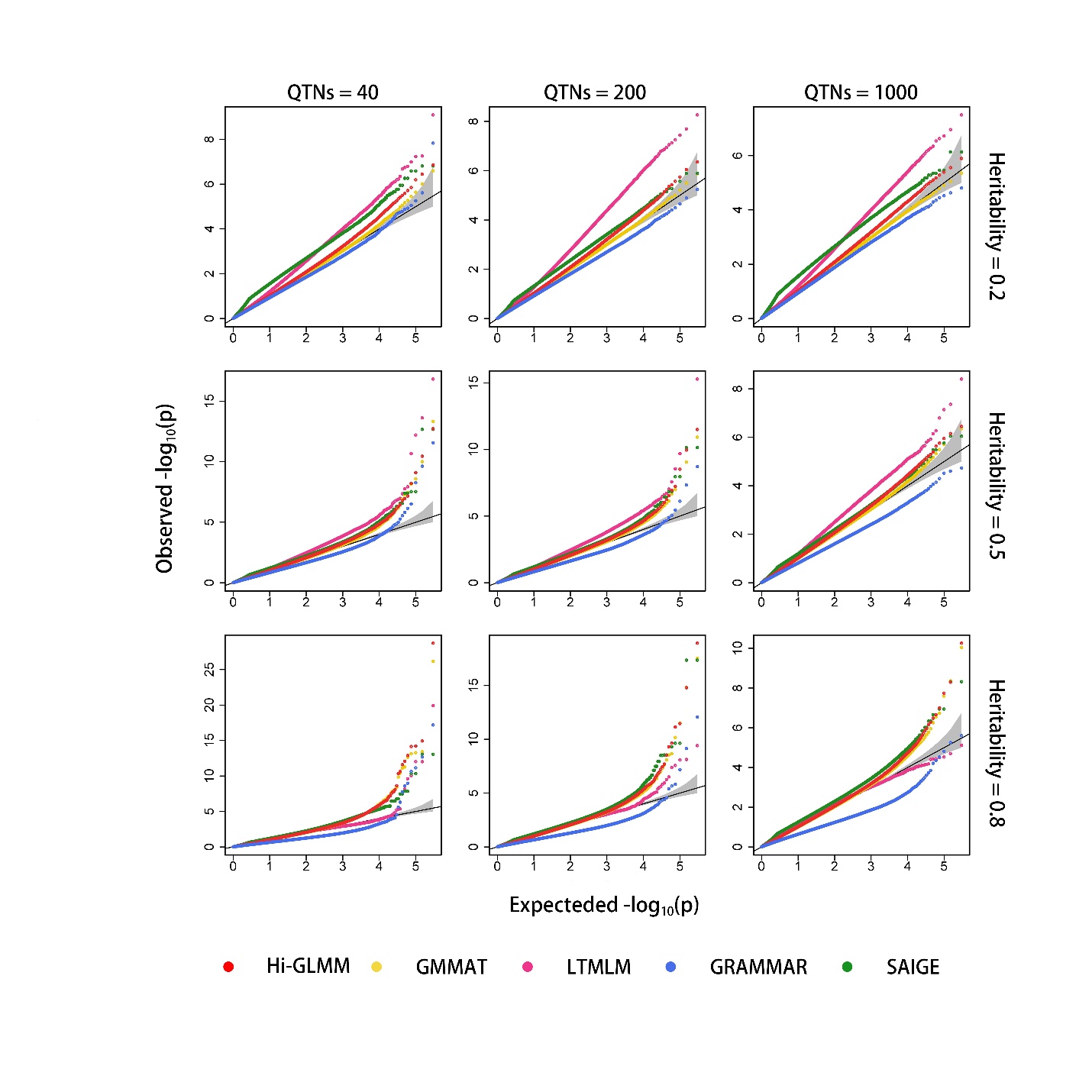

(2)

**Figure 2S:** Comparison in the ROC profiles between Hi-GLMM and the four competing methods. The ROC profiles are plotted using the statistical powers to detect QTNs relative to the given series of Type I errors. Here, the simulated phenotypes are controlled by 40, 200, 1,000 QTNs with the low, moderate and high heritabilities in 1) human and 2) maize.

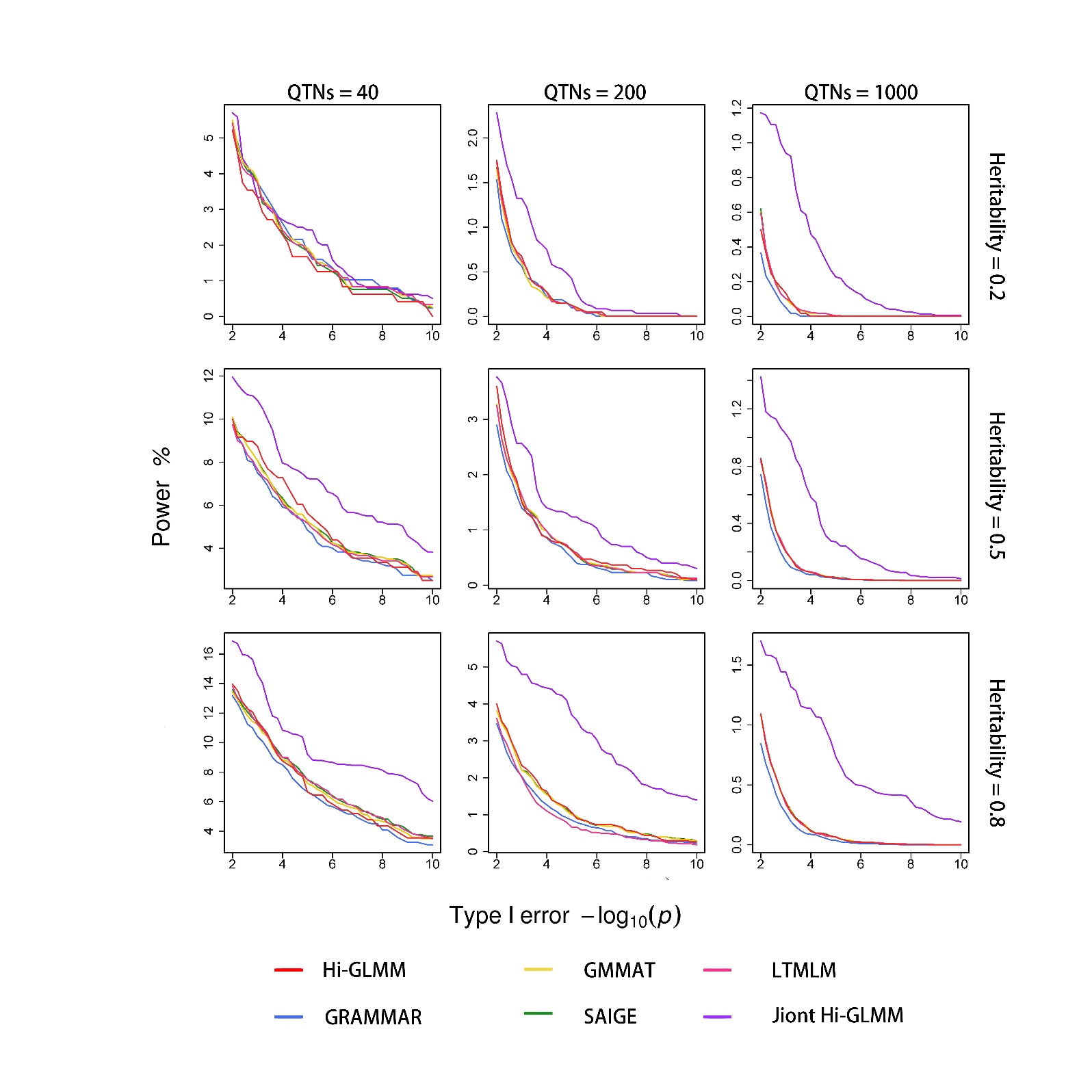

(1)

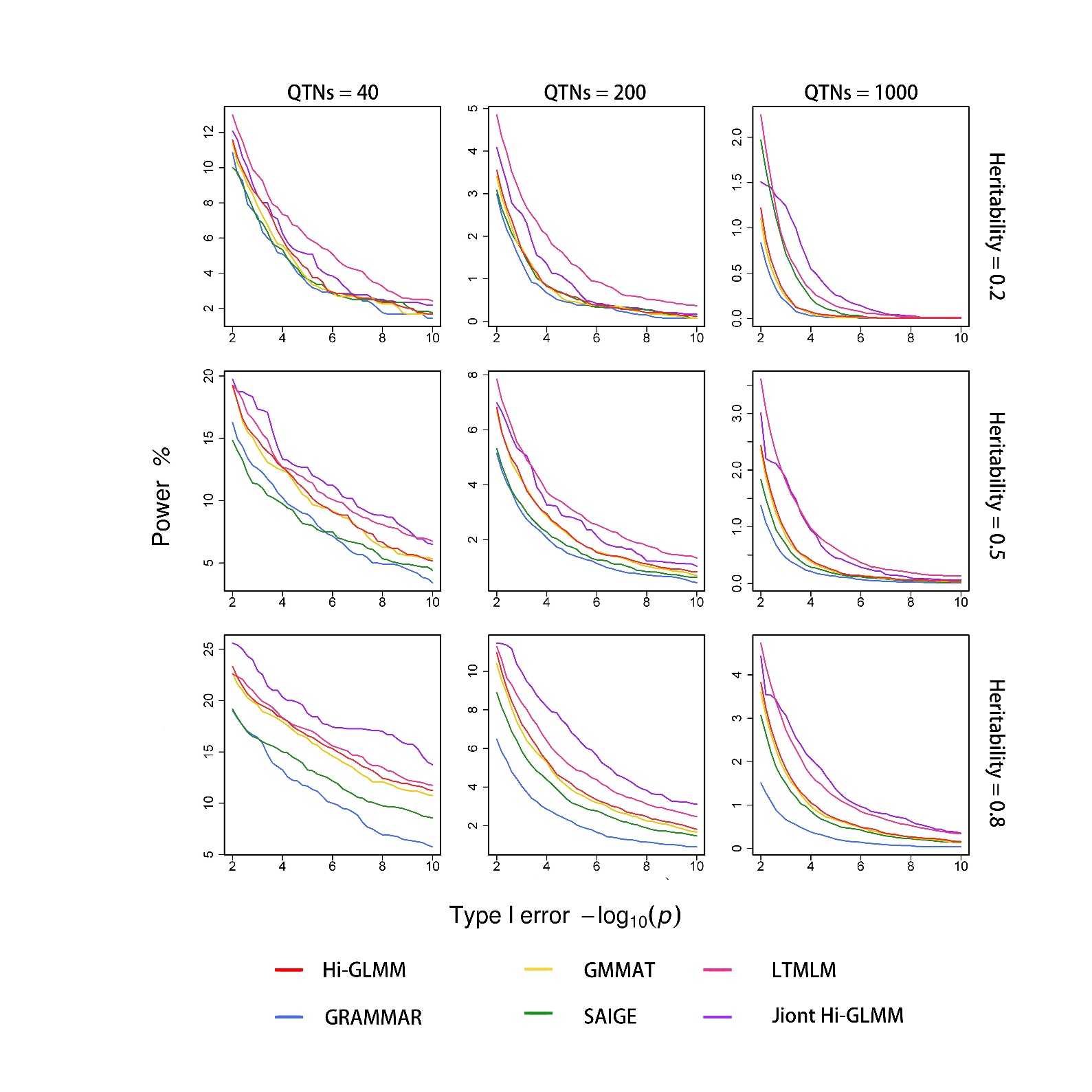

(2)

**Figure 3S:** Sensitivity of the Q-Q profiles to estimate heritabilities or GBVs for Hi-GLMM. Both GRAMMAR and SAIGE do not detect any QTN with the simulated GBVs. The simulated phenotypes are controlled by 200 QTNs with the low, moderate and high heritabilities in human and maize.

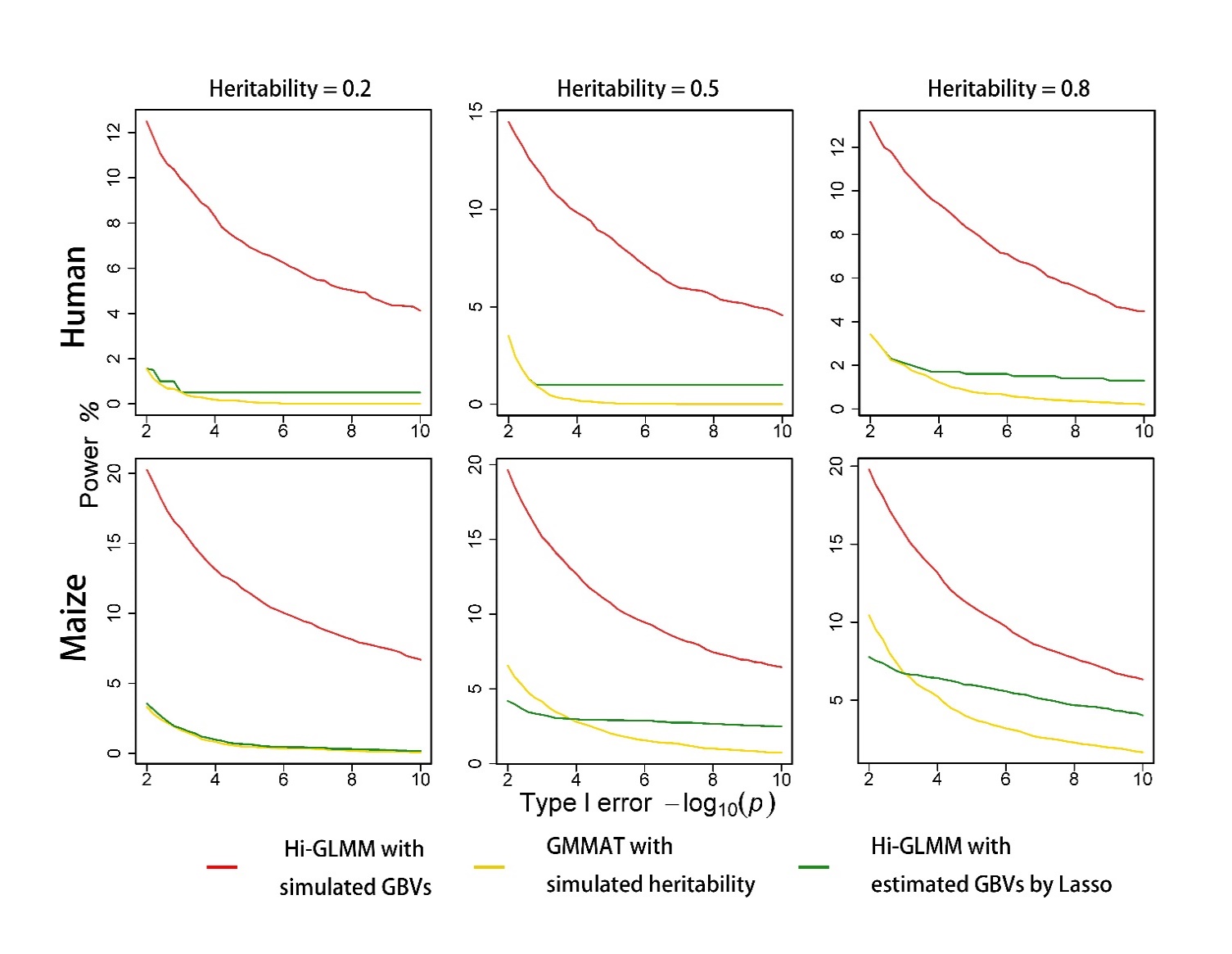

**Figure 4S**: Changes in Q-Q profiles with the number of sampling SNPs for Hi-GLMM. The simulated phenotypes are controlled by 40, 200 and 1,000 QTNs with the moderate heritability in human and maize.

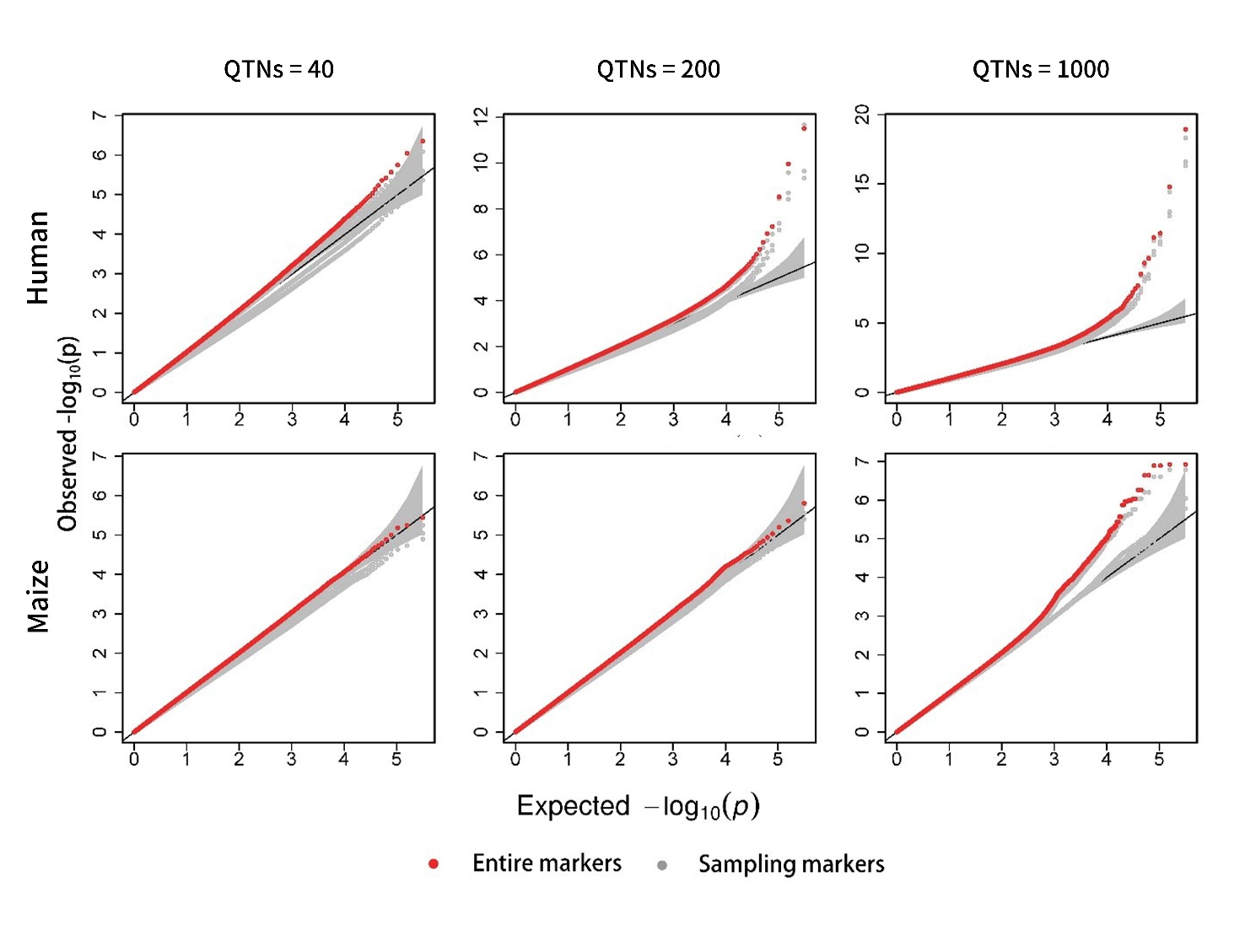

**Figure 5S:** Changes in ROC profiles with the number of sampling SNPs for Hi-GLMM. The simulated phenotypes are controlled by 40, 200 and 1,000 QTNs with the moderate heritability in human and maize.

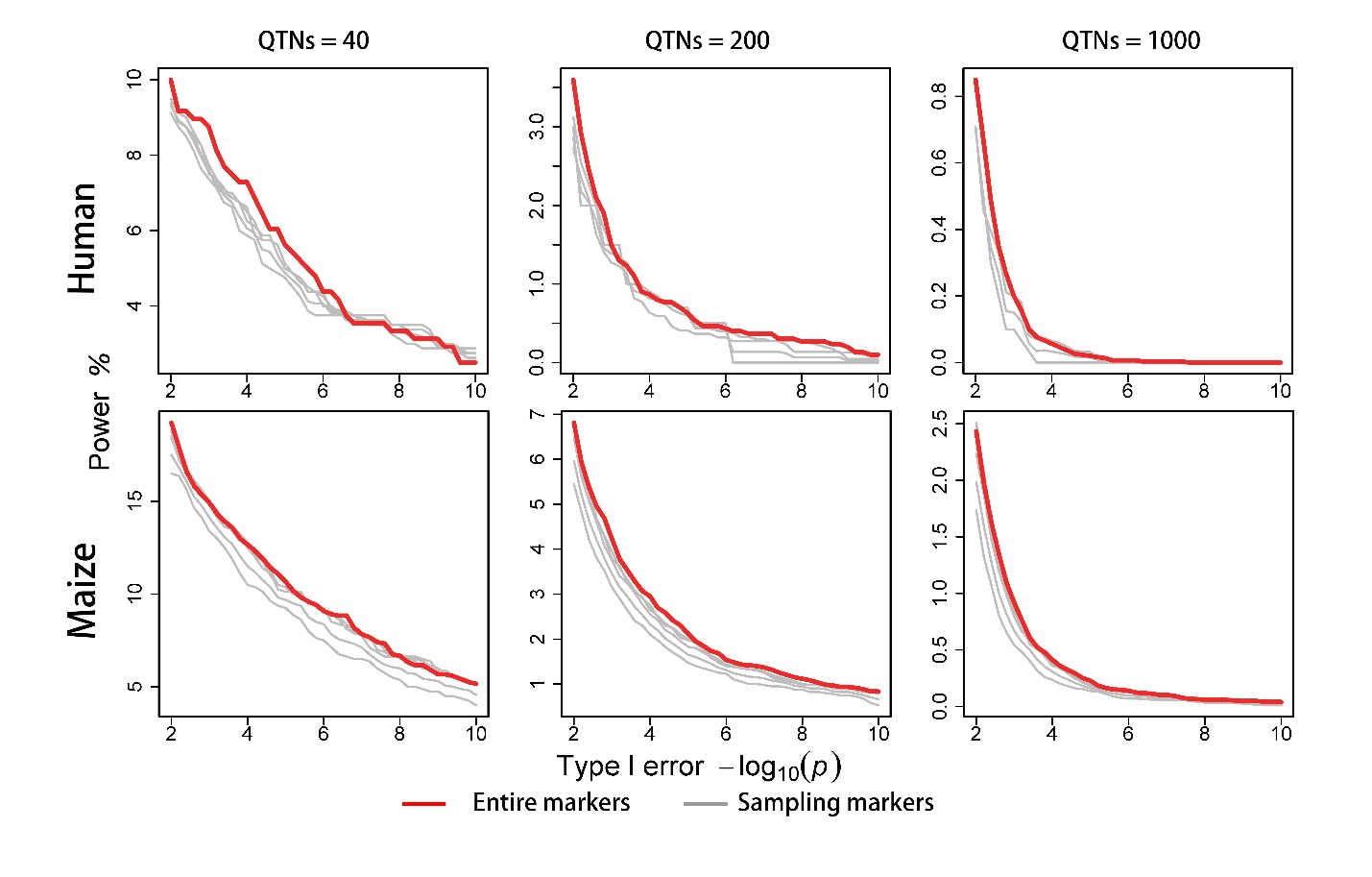

**Figure 6S:** Manhattan and Q-Q profiles for kernel colors in maize. The bottom is the Hi-GLMM with joint association analysis

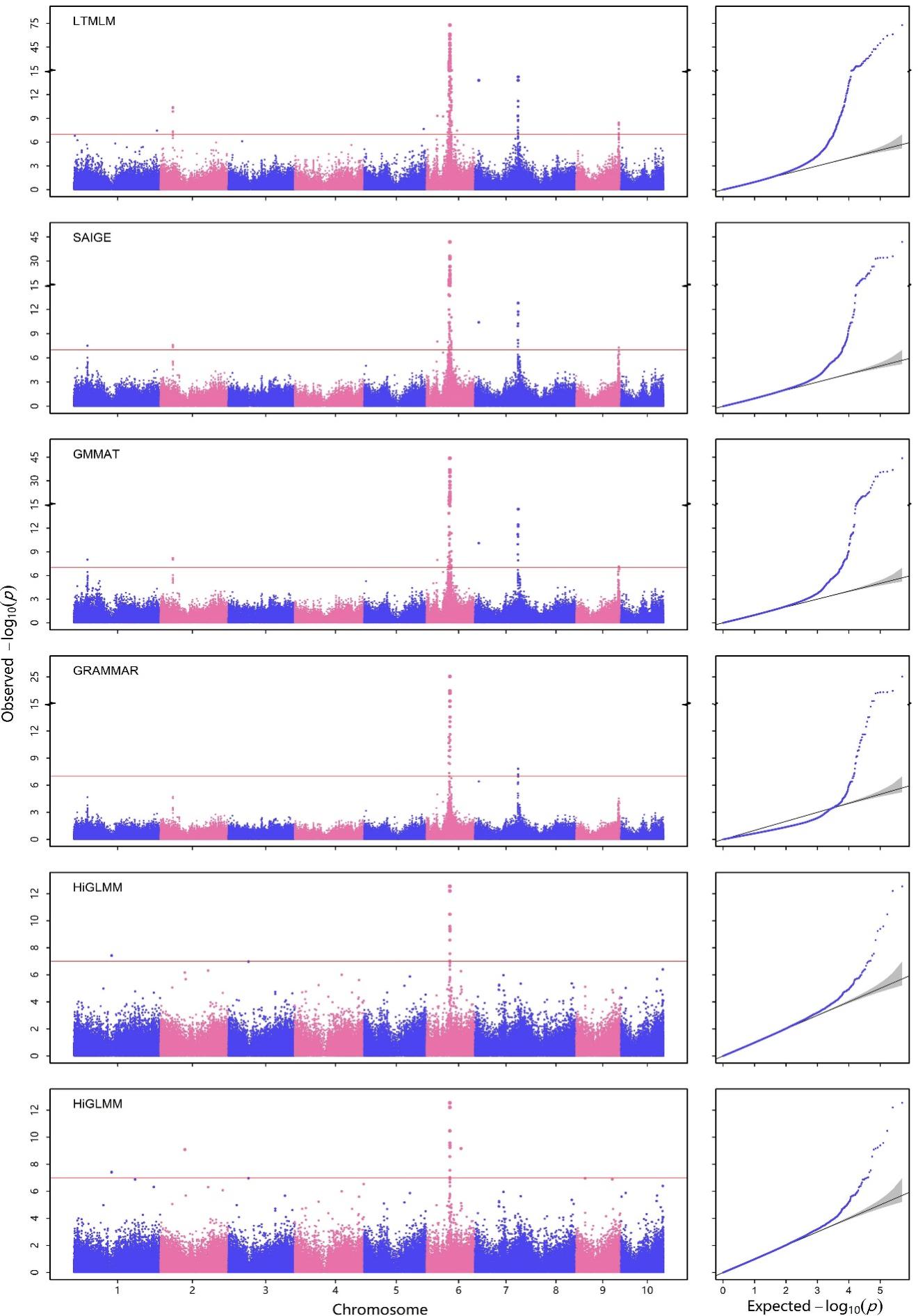

**Figure 7S:** Manhattan and Q-Q profiles for 1) BD; 2) CAD; 3) HT; 4) RA; 5) T1D and 6) T2D. The bottom is the Hi-GLMM with joint association analysis

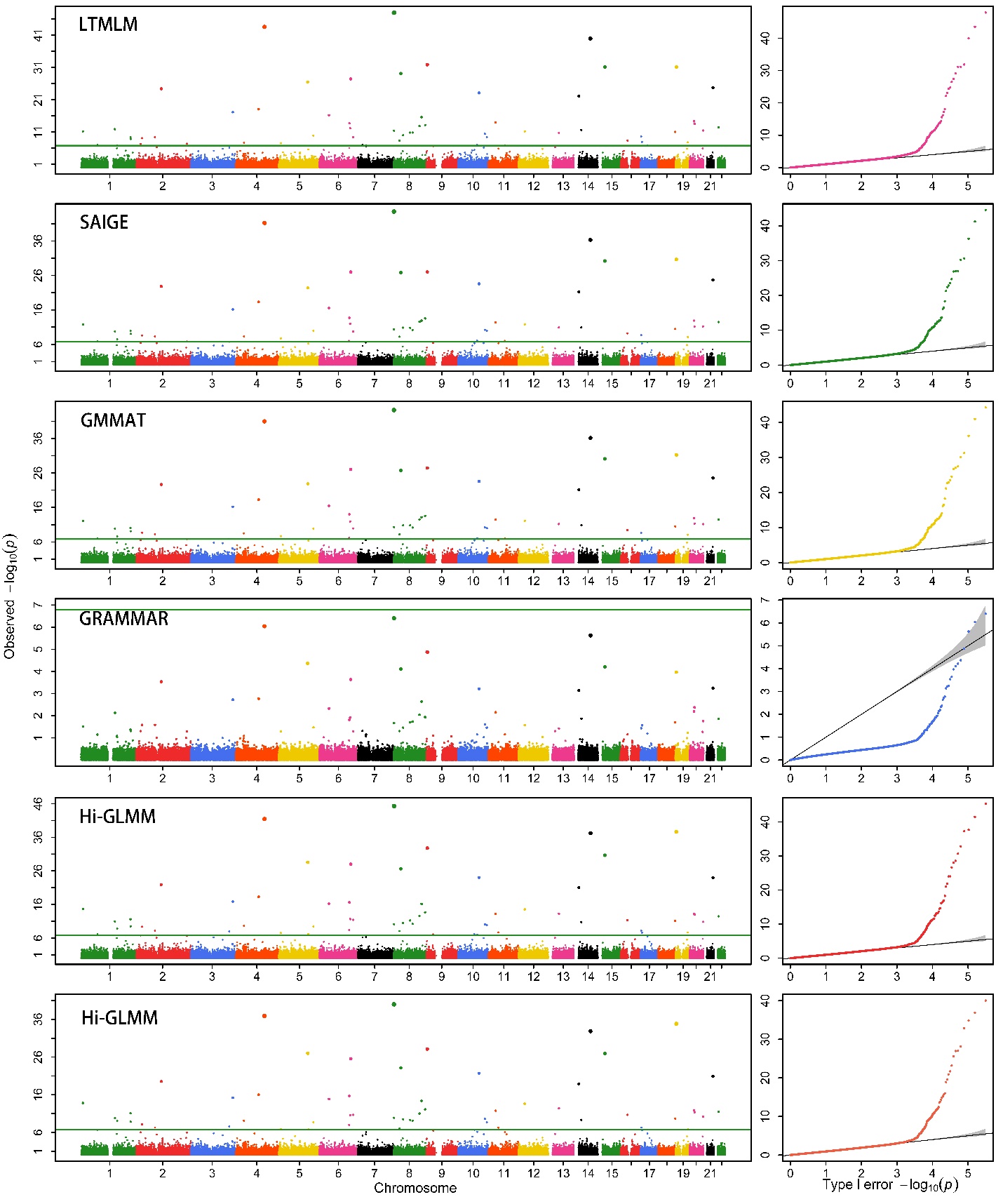

(1)

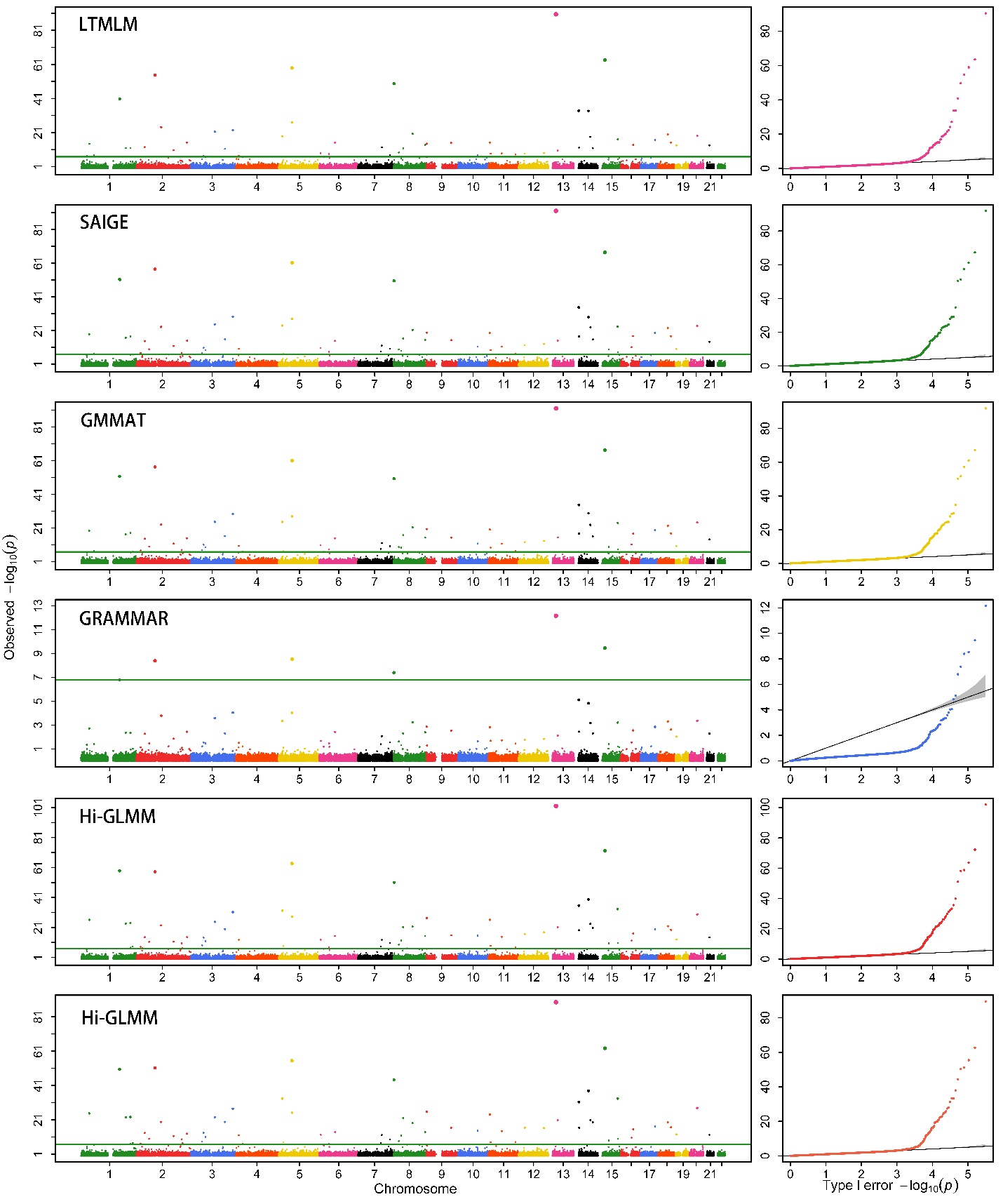

(2)

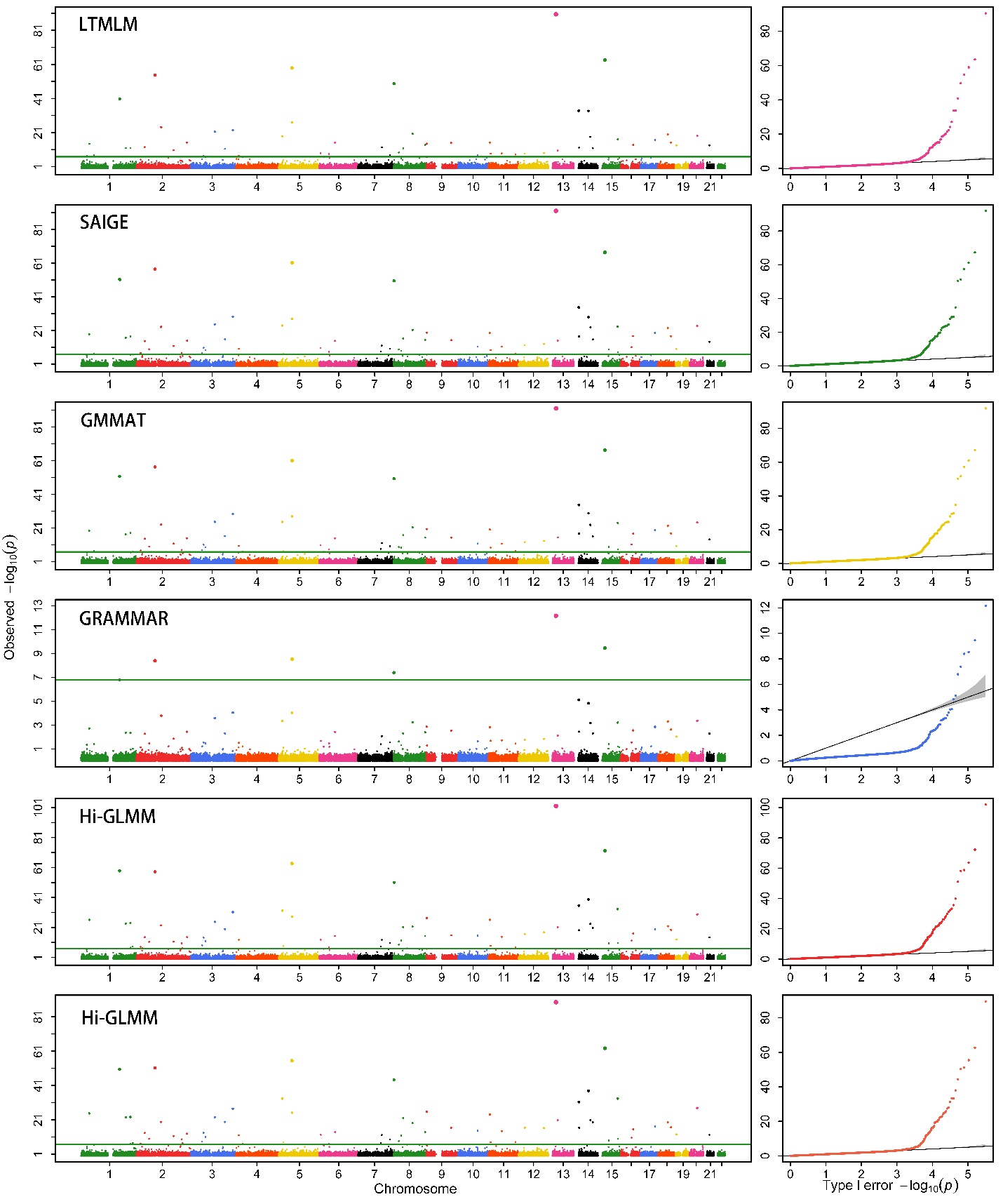

(3)

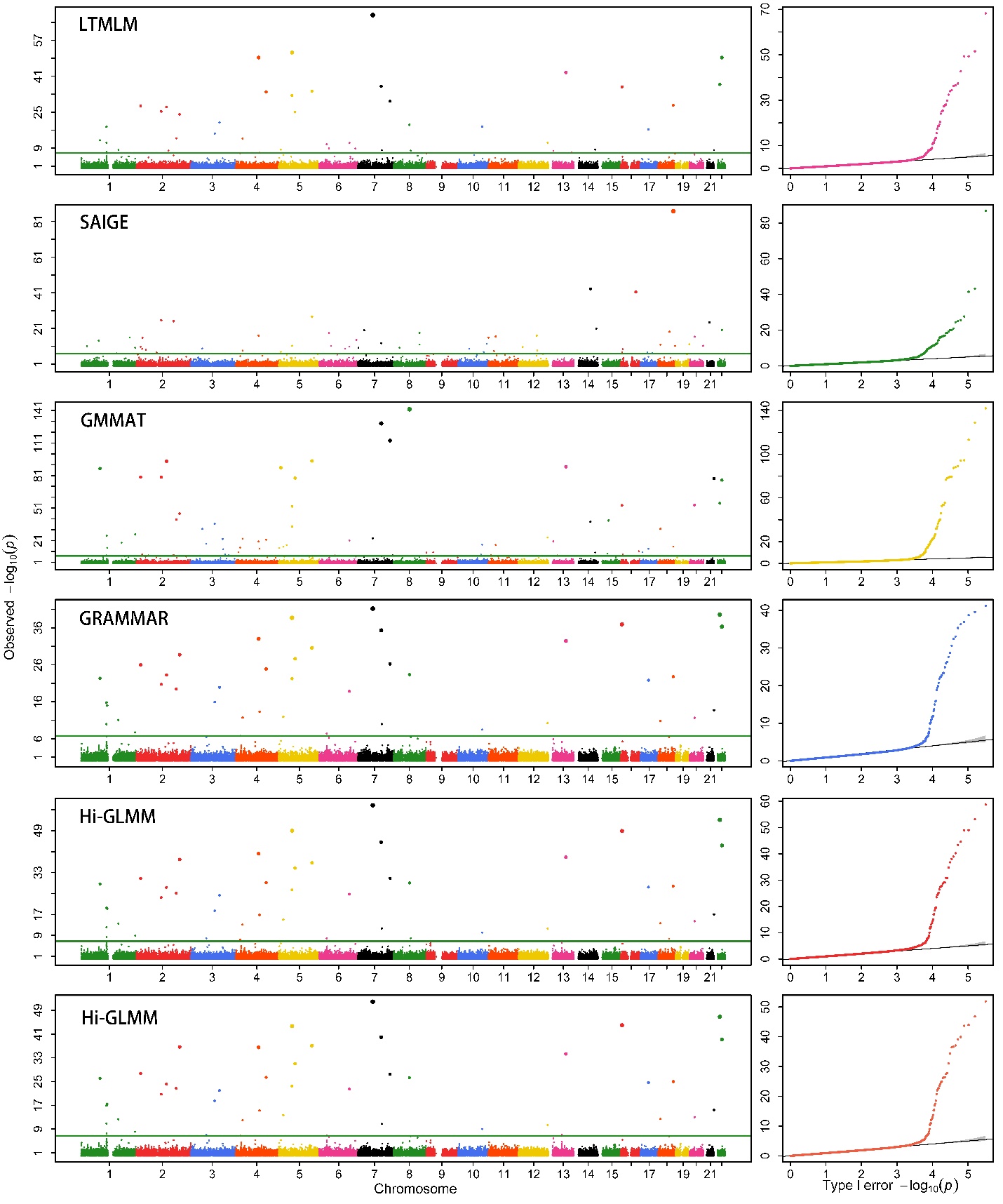

(4)

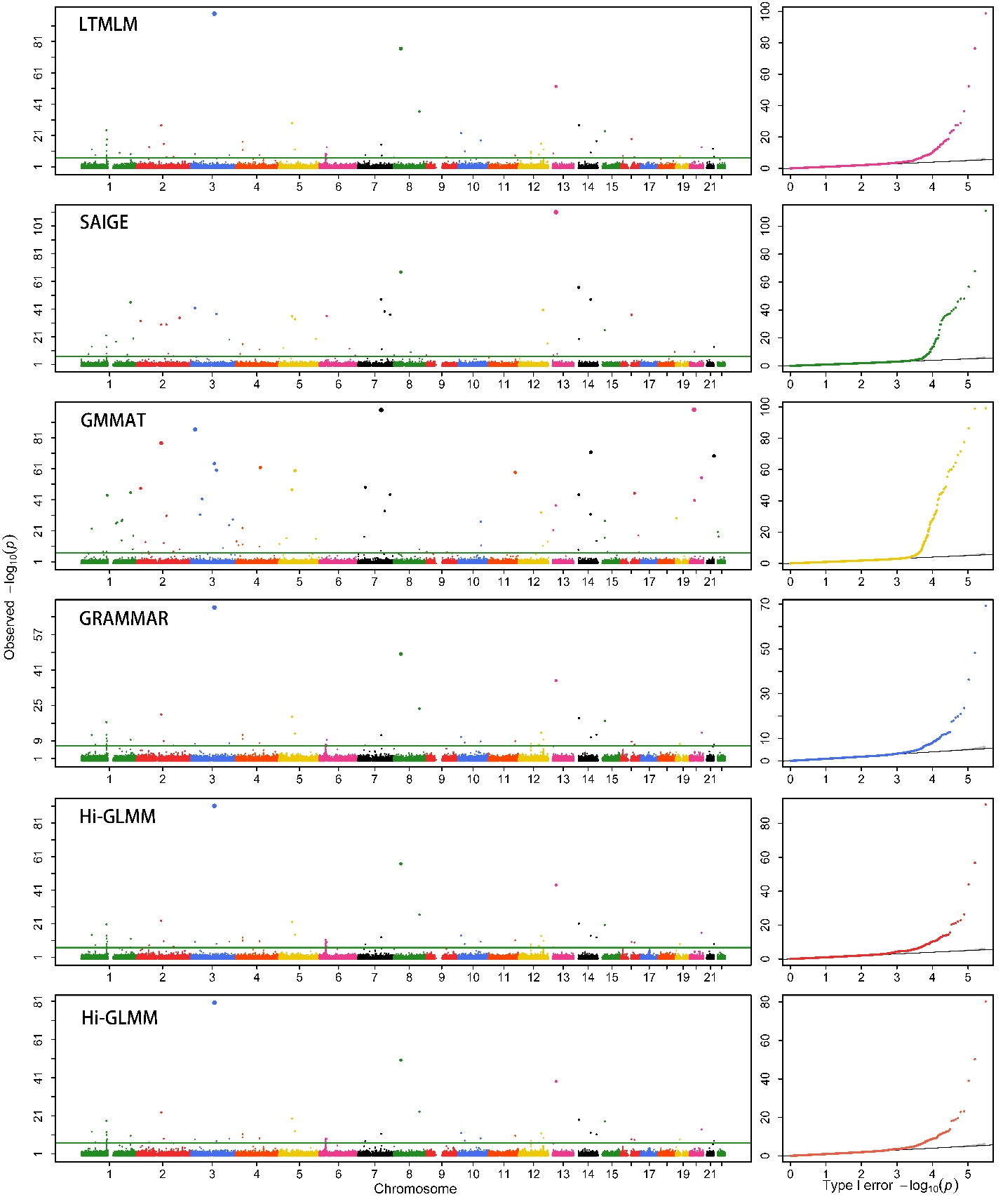

(5)

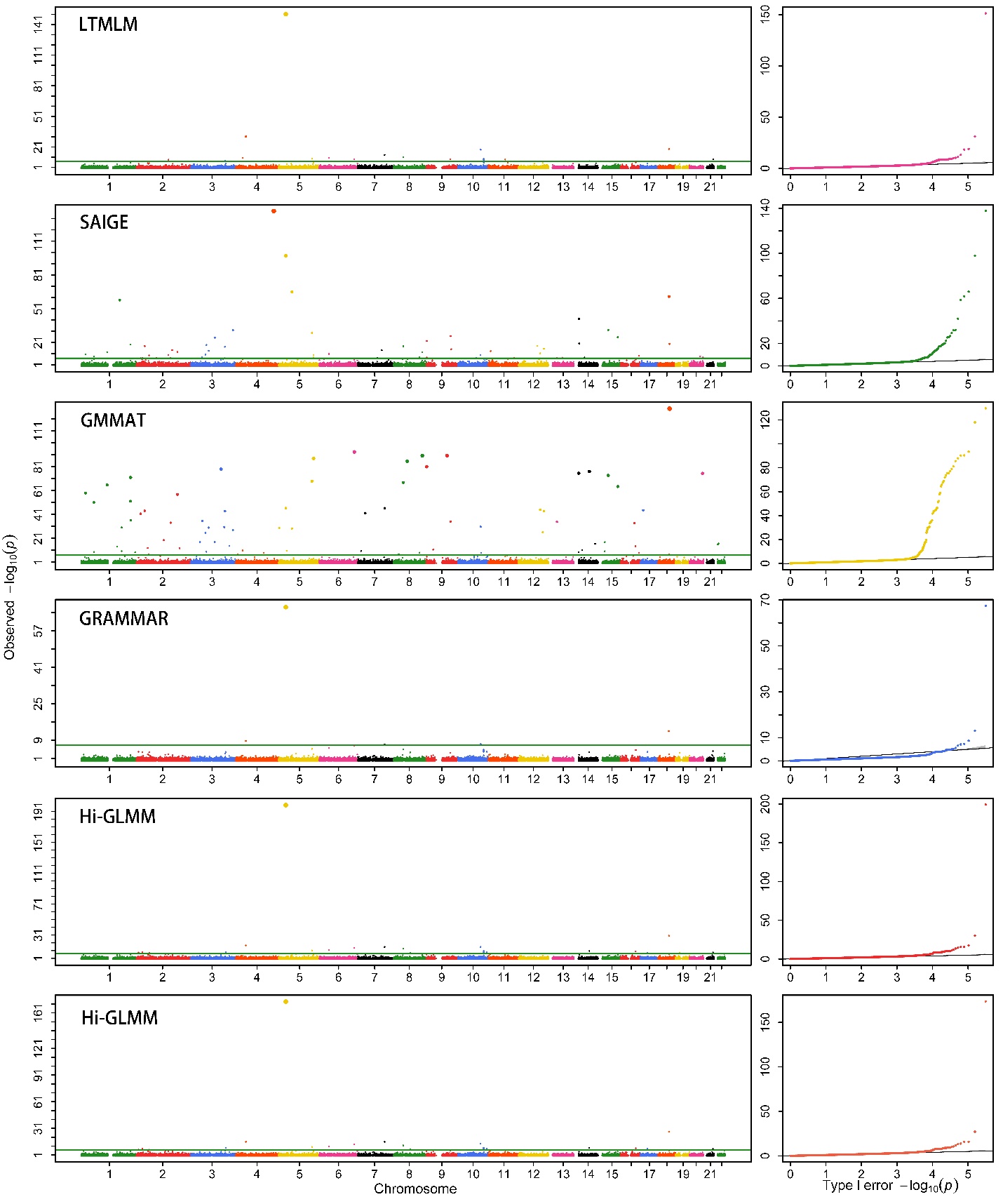

(6)

**Table 1S.** Genomic control values obtained with the five competing methods for all the simulated phenotypes

| Data | Method | 0.2 | | | 0.5 | | | 0.8 | | |
| --- | --- | --- | --- | --- | --- | --- | --- | --- | --- | --- |
|  |  | 40 | 200 | 1000 | 40 | 200 | 1000 | 40 | 200 | 1000 |
| Human | Hi-GLMM | 1(0.001) | 1(0.001) | 0.999(0.002) | 1.001(0.002) | 0.999(0.001) | 0.999(0.002) | 0.999 (0) | 0.999 (0) | 0.999(0) |
|  | GRAMMAR | 0.939(0) | 0.943(0) | 0.919(0) | 0.920(0.001) | 0.936(0.001) | 0.916(0) | 0.909(0.003) | 0.873(0) | 0.892(0) |
|  | LTMLM | 1(0.002) | 1.001(0.002) | 1.001(0.002) | 1(0.008) | 1.001(0.007) | 1.001(0.004) | 1(0.028) | 0.991(0.025) | 1.002(0.017) |
|  | GMMAT | 0.995(0) | 0.997(0) | 0.997(0) | 0.997(0) | 0.997(0) | 0.999(0) | 0.997(0) | 1.000(0) | 0.999(0) |
|  | SAIGE | 0.997(0.001) | 1(0.001) | 1(0) | 1(0.002) | 0.999(0.001) | 1.001(0) | 1(0.003) | 1.001(0.001) | 1.001(0) |
| Maize | Hi-GLMM | 1.002(0.006) | 1.005(0.007) | 1.007(0.009) | 1.005(0.006) | 1.003(0.009) | 1.014(0.008) | 1.005(0.044) | 1.003(0.021) | 1.014(0.003) |
|  | GRAMMAR | 0.907 (0.031) | 0.885 (0) | 0.917(0) | 0.781(1.686) | 0.745(0) | 0.999(0) | 0.531(0.001) | 0.556(0) | 0.529(0) |
|  | LTMLM | 1.576 (0.482) | 1.370(0.097) | 1.261(0.09) | 1.216(0.094) | 1.239(0.183) | 1.539(0.196) | 1.080 (0.52) | 1.144(0.499) | 1.112(0.258) |
|  | GMMAT | 0.998 (0.031) | 0.998(0.032) | 0.997(0) | 0.997(0.164) | 0.998(0) | 1.228(0) | 0.991 (0) | 0.995(0) | 0.997(0) |
|  | SAIGE | 1.956(0.480) | 1.536(0.481) | 1.975(0.480) | 1.368(0.685) | 1.335(0.522) | 1.331(0.480) | 1.343(0.580) | 1.366(0.455) | 1.423(0.458) |
